# An uncommon garden experiment: microenvironment has stronger influence on phenotypic variation than epigenetic memory in the clonal Lombardy poplar

**DOI:** 10.1101/2022.03.22.485169

**Authors:** Bárbara Díez Rodríguez, Cristian Peña, Paloma Pérez Bello, Julius Bette, Lena Lerbs, Tabea Mackenbach, Sven Wulle, Emanuele De Paoli, Koen J.F. Verhoeven, Katrin Heer, Lars Opgenoorth

**Affiliations:** Department of Biology, Philipps-University Marburg, Karl-von-Frisch Strasse 8 | D-35043 Marburg; Biodiversity and Conservation Biology, Swiss Federal Research Institute WSL, Zürcherstrasse 111 | CH-8903 Birmensdorf; Department of forest genetics, Albert-Ludwigs-Universität Freiburg, Bertoldstraße 17, 79098 Freiburg i. Br., Germany; Department of Agri-Food, Environmental and Animal Sciences, University of Udine. via delle Scienze 206, 33100 Udine, Italy; Department of Terrestrial Ecology, Netherlands Institute of Ecology (NIOO-KNAW), Droevendaalsesteeg 10, 6708 PB Wageningen; IGA Technology Services Srl. Via Jacopo Linussio 51, 33100 Udine UD, Italy

**Author notes:** Corresponding author: Katrin Heer.

**Keywords:** Phenotypic variation, *Populus nigra*, parental effects, epigenetic memory

## Abstract

Environmental changes can trigger phenotypic variation in plants through epigenetic mechanisms, but strong genetic influences on epigenetic variation and phenotypes make it difficult to isolate and study these effects. We investigated phenotypic plasticity using the Lombardy poplar *(Populus nigra* cv. ‘Italica’ Duroi), a globaly distributed clonal tree. We surveyed 14 functional traits related to tree growth, ecophysiological and phenological processes in poplar ramets collected along a wide geographical range in Europe and planted under common garden conditions. We investigated whether phenotypic variation was related to geography and historical bioclimatic data of the ramets’ sites of origin using linear mixed effect models. We found significant differences in among ramets from different geographic origins in tree height, number of stems per ramet and duration of bud flush. However, microenvironmental variation in the common garden, captured via block effects, had an even bigger impact on phenotypic variation than the environmental conditions at the sites of origin. Our results show that phenotypic variation in the ramets might be associated to the climate origin from different climates, suggesting possible epigenetic memory. However, such legacy effects might be quickly outweighed by new environmental conditions.

## 1. Introduction

Recent climatic extremes have shown that climate change already has severe impacts on temperate tree populations (Vitasse et al. 2019; Schuldt et al., 2020) and many studies suggest that due to their longevity trees are not able to adapt rapidly enough to keep pace with global climate change (Aitken et al., 2008; Bisbing et al., 2021). However, three factors have been suggested that might increase phenotypic variability and thus potentially the resilience of tree populations: intraspecific genetic variability (Benito Garzón et al., 2011; Pfenninger et al. 2021), (micro-)environmental variation (Scotti et al., 2016; Slavov et al., 2010; Sork et al., 2013), and epigenetic acclimation (Richards et al., 2017, Sow et al. 2018; Guarino et al.; 2015). For example, several studies on various *Populus sp*. genotypes have shown that patterns of phenotypic variation observed under common garden conditions usually follow latitudinal clines (Luquez et al., 2008; Howe et al., 2000; Ma et al., 2010; Rohde et al., 2010; McKown et al., 2013).

Phenotypic variation associated to clinal gradients has been observed in multiple functional traits, such as stomatal density (Gornall & Guy, 2008), specific leaf area (De Frenne et al, 2013), herbivory damage and herbivore abundance (Schemske, 2009; Robinson et al., 2012), stem height, relative growth rate, and total phenolic content (Luquez et al., 2008;). Furthermore, many of the these traits tend to show high heritability values in *Populus* species, suggesting that phenotypic differences have a major genetic component (Howe et al., 2000; McKown et al., 2014). Several factors that act a selective forces can contribute to this clinal variation, including temperature, precipitation, soil nutrient availability, and biotic agents (De Frenne *et al*., 2013). However, all of these studies dealt with genetically diverse (source) populations and since this genetic variability interacts with or might overshadow the impact of the microenvironmental and epigenetic marks on the phenotype, it is difficult to disentangle them in long lived organisms and thus quantify their relevance in structuring phenotypic variability. Here, we collected clonal Lombardy Poplar individuals across Europe and transferred them to a common garden environment. The genetic uniformity of this clone allows pinpointing epigenetic effects on phenotypic variation which can be induced by microenvironmental and large scale environmental variation.

Epigenetic modifications are chemical modifications in the DNA (for example, DNA methylation) that influence gene expression or function without altering the underlying DNA sequence (Bossdorf et al., 2007). These modifications can arise spontaneously or can be triggered by environmental conditions, and might be transmitted to the offspring (Latzel & Klimešová, 2010; Verhoeven & Preite, 2014; Munzbergova & Hadincova, 2016). If environment induces epigenetic marks, epigenetic variation can lead to “epigenetic memory” (Latzel et al., 2016). Clones thus offer a unique system to study epigenetically mediated plasticity (Latzel & Klimešová, 2010; Richards et al, 2017; Heer et al., 2018). First, because clones are characterized by low to zero genetic diversity, effects of epigenetic variation on trait variation will not be confounded with effects of genetic variation. Second, since clonal reproduction circumvents the epigenetic resetting associated with meiosis, epigenetic marks might be stable between clonal generations (Verhoeven and Preite, 2014).

Within single poplar genotypes, in absence of genetic effects on trait variation, phenotypic variation among plants can also arise based on transmission of parental environmental effects, potentially mediated by epigenetic mechanisms (Raj et al 2011). In our study, we worked with the so-called Lombardy poplar *(Populus nigra* cv. ‘Italica*’* Duroi) which is probably the widest distributed tree clone globally (CABI, 2022). The cultivar likely originated in the 18th century from one single male mutant tree of *P. nigra* located in central Asia (Elwes and Henry, 1913) and its cuttings were introduced to Italy, from where its cuttings were distributed worldwide for ornamental purposes and as a source of timber (Wood, 1994). It is assumed that almost all Lombardy poplars are the result of artificial propagation performed by humans. This unique origin and the wide geographical distribution of the clone makes the cultivar a perfect study system to investigate epigenetically induced phenotypic variation in a long-lived plant species and its potential role in plant adaptation (Vanden Broeck et al., 2018).

In this study, we aimed to investigate the effect of previous environments on the phenotypic variation of the next clonal generation of Lombardy poplar ramets growing under common garden conditions. We surveyed 14 functional traits related to tree growth, ecophysiological and phenological processes under common garden conditions in one to two growing seasons. We genotyped all ramets established in the garden to determine clonal identity. Using historical bioclimatic data from each region where the ramets were collected, we related phenotypic variation to geographic and climatic gradients. We hypothesized that (1) phenotypic variation in functional traits would correlate with macro-climatic gradients from the ramets sites of origin and (2) that microenvironmental differences in the common garden would not have any effect on the clone phenotypes.

## 2. Material and methods

### 2.1. Plant material and common garden design

Between February and March 2018, cuttings from *Populus nigra cv* ‘Italica’ clones were collected in Europe across geographical gradients that spanned from 41° to 60° N and −5° to 25° E approximately (Figure 1).

**Figure 1.**
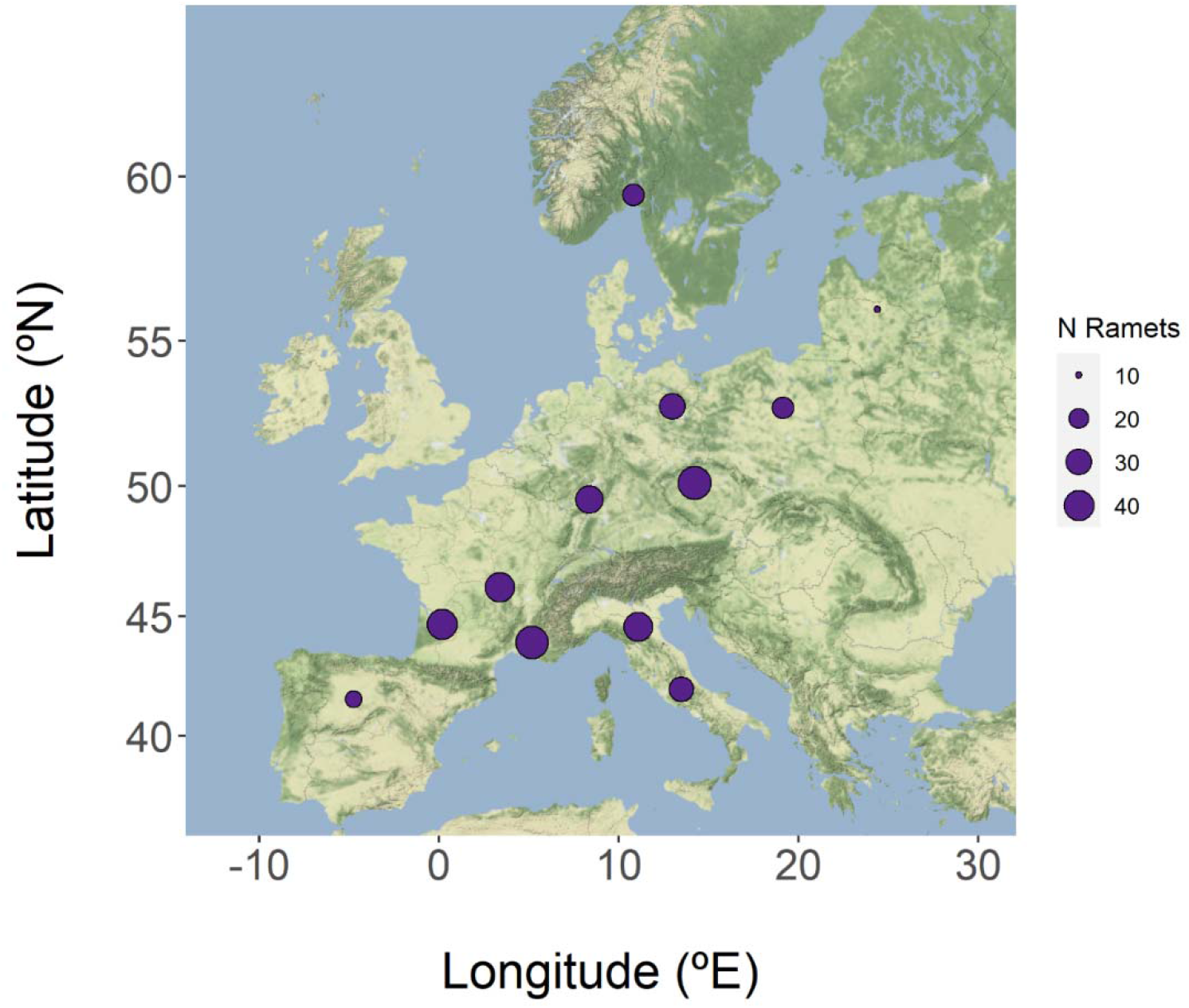
Distribution of sample sites of Lombardy poplars and number of ramets collected in each location.

**Figure 2.**
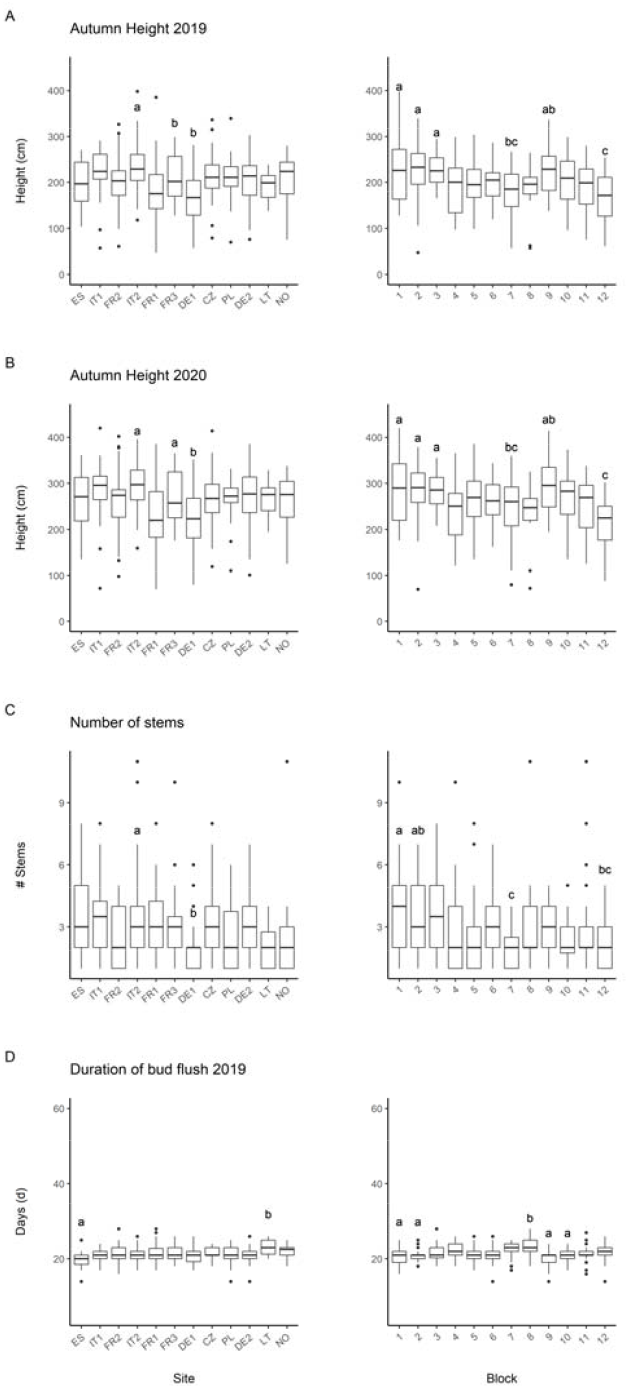
Phenotypic variation in traits for which significant differences among geographic origins were found. Boxplots are ordered by site (left column) or by common garden block (right column). Sites are ordered from south to north according to their geographic coordinates and labelled by the sample site code (ISO 3166 standard country code): ES: Spain; IT1: Italy 1; FR2: France 2; IT2: Italy 2; FR1: France 1; FR3: France 3; DE1: Germany 1; CZ: Czech Republic; PL: Poland; DE2: Germany 2; LT: Lithuania; NO: Norway. Sites or blocks that are significantly different (p < 0.05) are labelled with different letters.

To cover a wide bioclimatic range, twelve sampling sites were selected that covered seven different Köppen-Geiger climate subtypes (Peel et al., 2007). At each site, cuttings of approximately 30 cm in length were sampled from 50 to 56 different individuals within a 25 km radius, except for the sites in Spain, Poland, and Lithuania, where only 24, 27, and 12 individual trees were found within the sampling radius, respectively. Source trees (ortets hereafter) were tagged and georeferenced. During the first week of May 2018, the cuttings (ramets hereafter) were planted on a previously tilled lawn in the Marburg Botanical Garden (Germany). The common garden is located at 50º 48’ 02.7” N, 8º 48’ 24.8” E, at an elevation of 328 m within the Cfb (temperate oceanic climate) of the Köppe-Geiger climate classification. To control weeds, the trial area was covered with water-permeable plastic foil (Agrolys BL100 #25/12.5, Beaulieu Technical Textiles) with openings of about 10 cm x 10 cm placed over the ramets. Weeds that grew through the openings in the foil were controlled manually if necessary. The common garden area was not shaded in any way, allowing the ramets to grow under direct sunlight. No herbicides, pesticides, or fertilizers were used in the common garden. The area was fenced to exclude herbivory or other disturbances by deers and wild boars. The ramets were planted in a random block design (Figure S1) composed of 12 blocks in a 3 by 4 array with 40-45 cuttings per block. In each block, two cuttings from 1-5 individual ortets from each sampling site were planted. After one of the ramets had successfully established, the other was removed from the garden. The ramets were planted with a distance of 1 m between trees and were watered frequently until the end of summer. In total, 549 poplar ramets were planted, of which 433 survived the summer of 2018.

### 2.2. Genotyping of ramets

To determine whether the ramtes really belonged to a single clone line, leaf samples were collected and genomic DNA was extracted between July and August 2018. DNA samples were isolated with the PeqGOLD Plant DNA mini kit (PEQLAB Biotechnologie GmbH, Erlangen, Germany). The clones were genotyped at 5000 genomic loci, equally distributed across the 19 *P. nigra* chromosomes and selected among a larger set of polymorphic sites identified in Scaglione et al. (2019). Sequencing was carried out using the Allegro Targeted Genotyping protocol from NuGEN Technologies (TECAN) by IGATech (Udine, Italy), yielding information for 8,218 SNP positions. Among these, 3,313 SNPs (40.3%) were monomorphic across all trees, bringing the number of informative SNPs to 4,905. Three adult *P. nigra cv* ‘Italica’ clones from the botanical garden in Marburg were included in the sequencing design as controls. Of the 433 ramets established in the common garden, 374 belonged to a single genotype with a mean number of pairwise differences among individual ramets (including the adult clones of the botanical garden) equal to 96 ± 40 (s.d) SNPs, corresponding to 1.2% of all SNP positions analyzed (Supplementary Figure S2). We considered these trees as *bona fide* Lombardy poplars and focused the rest of the study on this subset.

### 2.3. Trait measurement in the common garden

To study phenotypic variation across geographic and climatic gradients, 14 phenotypic traits were measured under common garden conditions. These were divided into five categories: tree growth, ecophysiology, biotic stress damage, leaf chemistry, and phenology.

#### 2.3.1. Tree growth traits

Ramet diameters were measured with an electronic caliper before the ramets were planted in 2018. Tree height was measured at the ground level at the end of the growing season in 2018, 2019, and 2020, after all ramets had completed bud set. The number of stems was counted at the end of the 2018 growing period and corresponds to the number of initial sprouts that appeared in the ramets in the summer. The active growth rate was calculated as the ratio between the height gain (2018 to 2019) and the number of days of growth. The growth period was calculated as the number of days between the last stage of bud flush (Stage 5 in Azad, 2012) and the first stage of bud set (Stage 2.5 in Rohde et al., 2011, see detailed description below).

#### 2.3.2. Ecophysiological traits

##### i) Leaf traits

In July 2020, leaves from a subset of 163 ramets randomly chosen were sampled for assessing specific leaf area (SLA) and leaf mass per area (LMA). Three branches from each ramet were chosen randomly. In poplar, new leaves are produced from the apical shoot, so to measure a fully mature leaf, we collected the eighth leaf counting from the first fully unfolded leaf in the apical shoot of the branch. Only healthy leaves with no signs of biotic or abiotic stress were sampled. If the eighth leaf did not meet the criteria to be considered undamaged, the next healthy leaf was sampled. To assess leaf area, leaves were scanned on a flat white background at 300 dpi using an Epson perfection V370 photo scanner. Based on these scans, leaf area was calculated using the WinFOLIA leaf area analysis software (Regent Instruments Inc.). Leave dry weight was determined by drying leaf samples in an oven at 60° C for 2 days and weighted with a balance precision of 0.001 g. SLA was calculated as the ratio between leaf area and leaf dry weight and LMA as 1/SLA.

##### ii) Stomatal density

Immediately after the leaves were scanned, two of the three leaves were randomly chosen to determine stomatal density (SD). Epidermal impressions of the abaxial side of the leaves were obtained by creating an imprint of the leaf surface with clear fingernail polish (Maybelline Superstay 7 Days Gel Nail Color). The polish imprints were mounted in permanent microscope slides with glass covers and photographed under a Zeiss Axio Lab.A1 microscope at 10x magnification in a 450 × 350 μm field. The number of stomata was counted using the StomataCounter software (Fetter et al., 2019). The software automatically annotates visible stomata. In addition, we visually double checked the automated annotation and added undetected stomata manually.

#### 2.3.3. Biotic Stress traits

Herbivory damage was assessed in July 2019. A single randomly chosen branch in each ramet was selected and fifty leaves in the branch were scored for the presence or absence of herbivore damage. The number of leaves was counted from the bottom of the branch, and the seven leaves around the shoot area were excluded from the scoring system. When the ramet was too small to have fifty leaves in a single branch, a lower number of leaves was selected, except for two ramets that were completely excluded from the surveys due to their small size. The percentage of damage was calculated as the ratio of the number of damaged leaves to the number of undamaged leaves.

#### 2.3.4. Leaf chemistry traits

In June 2020 leaf anthocyanin content (Anth), chlorophyll content (Chl), flavonol content (Flav), and Nitrogen Balanced Index (NBI, a plant status indicator directly correlated with massic nitrogen content) were estimated using a Dualex Scientific meter (FORCE-A, Orsay, France) The meter measures the light transmittance ratio at two different wavelengths in the near-infrared and far-red range to calculate the chlorophyll content. Flavonol and anthocyanin content is calculated based on the amount of light absorbed by polyphenols and the amount that reaches the chlorophyll in the mesophyll. The same subset of clones used for the SLA analysis was used for leaf chemistry analysis. Ten mature healthy leaves per clone were randomly chosen. The Dualex meter measurements were taken on the center of the adaxial side of the leaves next to the midrib.

#### 2.3.5. Phenological traits

Bud phenology, specifically bud set and bud flush, was monitored by focusing on the main apical bud. Bud set was scored in autumn 2018 and 2019 according to the scoring scale designed by Rohde et al. (2011), and bud flush was scored in spring 2019 and 2020 using the scale suggested by Azad (2012). The bud set scale spans seven stages, from stage 3 (apical shoot fully growing) to stage 0 (bud set), while the bud flush scale consists of six stages, from stage 0 (dormant bud) to stage 5 (leaves fully unfolded). Bud stages were recorded every two to three days at the beginning of the monitoring period and then daily until the apical buds of all clones had reached the final stage of their respective scales. Because bud phenology in *P. nigra* depends greatly on day length and temperature and all ramets were exposed to the same cues, no variation in the day when bud set or bud flush started was expected. The duration of bud formation, however, has been shown to differ in identical genotypes growing under different conditions (Rohde et al., 2011). Therefore, phenological traits were defined as duration of bud set or duration of bud flush, which equaled the number of days it took each ramet to reach from the first stage to the last stage, respectively.

### 2.4. Climatic variables and climatic gradients

Climatic data for each of the locations of the ortets were obtained from the CHELSA time-series data set (Karger et al., 2017). The CHELSA data set covers the period between 1979 and 2013 and provides gridded data at a resolution of 30 arcsec (∼ 1km). A Principal Component Analysis (PCA) was performed using all bioclimatic variables (bioclims, BIO 1-19). Individual coordinates for PC1 and PC2 were obtained and included as variables that represented climatic gradients. The bioclims that contributed the most to PC1 were all related to temperature variables (except for BIO 19, precipitation of the coldest quarter), while the most contributing bioclims in PC2 were related to precipitation variables (except for BIO 2, mean diurnal range). All bioclims and their contributions to each PC are described in Table S1.

### 2.5. Statistical analysis

All statistical analyses were performed in R (version 4.0.3; R core team, 2020). Basic descriptive statistics were calculated for all traits (Table 1). To assess if variation in cutting diameter and developmental processes such as the number of stems produced might have an effect in other phenotypic traits, all traits where correlated with each other using Pearson’s Product-moment Correlation with the *cor* function from the base R *stats* package. The effects of geographic origin and microenvironmental conditions on phenotypic variation were tested using linear mixed-effects models (LMMs). Phenotypic traits were the response variable in all models. Two models were fitted for each trait, one with bioclimatic variables (PC1 and PC2) and one with sampling site as fixed effects. To disentangle the possible effects of microenvironmental conditions in the common garden from the effects the ortet provenances had on phenotypic variation, the garden block in which the ramets were planted was included as a random effect in all the LMMs (Table 2). Since many of the traits analyzed were correlated with tree height and cutting diameter, this source of variation was accounted for by also including these variables in the models as fixed effects. Several models were fitted including tree height in 2018 or 2019, cutting diameter or both variables as fixed effects. Based on the best marginal R^2^ values, which we calculated with the *rsquared* function (R package *piecewiseSEM*, version 2.1.2) we decided which variable was included in the model. The LMMs were fit using the *lmer* function from the *lme4* package (version 1.1-23; Bates et al., 2015). Differences between groups were tested using the *emmeans* R package (version 1.6.3; Lenth, 2021) and p-values were corrected for multiple pairwise comparisons using the Bonferroni correction.

**Table 1.** Phenotypic traits of Lombardy poplar ramets from across Europe under common garden conditions.

| Phenotypic trait | Year | N | Min | Max | Median | Mean ± SD |
| --- | --- | --- | --- | --- | --- | --- |
| Tree growth |  |  |  |  |  |  |
| Cutting diameter (cm) | 2018 | 373 | 0.48 | 4.35 | 2.27 | 2.43 ± 0.70 |
| Growth rate (cm day<sup>-1</sup>) | 2019 | 342 | 0.08 | 1.76 | 0.83 | 0.84 ± 0.18 |
| Height Autumn (cm) | 2018 | 374 | 13.00 | 229.50 | 101.75 | 100.28 ± 39.16 |
| Height Autumn (cm) | 2019 | 372 | 47.00 | 398.00 | 207.50 | 203.16 ± 55.45 |
| Height Autumn (cm) | 2020 | 371 | 70 | 420 | 269 | 263.12 ± 64.11 |
| Stems (#) | 2018 | 374 | 1 | 11 | 2 | 2.95 ± 1.86 |
| Ecophysiology |  |  |  |  |  |  |
| LMA (mg mm<sup>-2</sup>) | 2020 | 163 | 0.04 | 0.10 | 0.06 | 0.06 ± 0.01 |
| SLA (mm<sup>2</sup> mg<sup>-1</sup>) | 2020 | 163 | 10.13 | 26.80 | 16.34 | 16.35 ± 2.06 |
| Stomatal Density (mm<sup>-2</sup>) | 2020 | 126 | 142.86 | 269.84 | 206.35 | 206.15 ± 28.71 |
| Biotic stress |  |  |  |  |  |  |
| Herbivory damage (%) | 2019 | 367 | 2.00 | 52.00 | 28.00 | 27.49 ± 9.64 |
| Leaf Chemistry |  |  |  |  |  |  |
| Anthocyanins (RAU) | 2020 | 155 | 0.12 | 0.18 | 0.14 | 0.14 ± 0.01 |
| Chlorophyll (μg cm<sup>-2</sup>) | 2020 | 164 | 24.79 | 35.38 | 30.28 | 30.18 ± 2.11 |
| Flavonols (RAU) | 2020 | 164 | 1.88 | 2.16 | 2.07 | 2.07 ± 0.05 |
| Nitrogen Balanced Index | 2020 | 164 | 11.83 | 17.70 | 14.67 | 14.62 ± 1.12 |
| Phenology |  |  |  |  |  |  |
| Budflush (d) | 2019 | 368 | 14 | 28 | 21 | 21.33 ± 2.26 |
| Budflush (d) | 2020 | 368 | 8 | 34 | 22 | 22.14 ± 5.88 |
| Budset (d) | 2018 | 374 | 12 | 68 | 22 | 22.63 ± 6.60 |
| Budset (d) | 2019 | 344 | 19 | 39 | 25 | 27.28 ± 4.42 |

**Table 2.** Climatic and geographic model coefficients

| Trait Class | Phenotypic trait | Model | Fixed effects |  |  | Model R<sup>2</sup> |  |
| --- | --- | --- | --- | --- | --- | --- | --- |
|  |  |  | Fixed 1 | Fixed 2 | Fixed 3 | R<sup>2</sup> marginal | R<sup>2</sup> conditional |
| Biomass | Height Autumn 2018 | Climatic | PC1 | PC2 | Cutting *** | 0.058 | 0.139 |
|  |  | Geographic | Site | Cutting * | - | 0.093 | 0.172 |
|  | Height Autumn 2019 | Climatic | PC1 | PC2 | Cutting *** | 0.061 | 0.170 |
|  |  | Geographic | Site * | Cutting | - | 0.102 | 0.206 |
|  | Height Autumn 2020 | Climatic | PC1 | PC2 | Cutting *** | 0.064 | 0.192 |
|  |  | Geographic | Site * | Cutting |  | 0.113 | 0.234 |
|  | Growth rate 2019 | Climatic | PC1 | PC2 * | Cutting ** | 0.036 | 0.116 |
|  |  | Geographic | Site | Cutting * | - | 0.060 | 0.144 |
|  | Number of stems | Climatic | PC1 * | PC2 | Cutting | 0.015 | 0.065 |
|  |  | Geographic | Site * | Cutting | - | 0.065 | 0.115 |
| Ecophysiology | Leaf Mass Area | Climatic | PC1 | PC2 | Height 19 * | 0.043 | 0.092 |
|  |  | Geographic | Site * | Height 19 |  | 0.083 | 0.121 |
|  | Specific Leaf Area | Climatic | PC1 | PC2 | Height 19 ** | 0.053 | 0.084 |
|  |  | Geographic | Site * | Height 19 * | - - | 0.098 | 0.121 |
|  | Stomata Density | Climatic | PC1 | PC2 | Height 19 | 0.030 | 0.139 |
|  |  | Geographic | Site | Height 19 | - | 0.077 | 0.168 |
| Biotic stress | Herbivory damage | Climatic | PC1 ** | PC2 | Height 19 * | 0.035 | 0.104 |
|  |  | Geographic | Site | Height 19 | - | 0.061 | 0.126 |

**Table 2** (Continued)
| Trait Class | Phenotypic trait | Model | Fixed effects |  |  | Model R<sup>2</sup> |  |
| --- | --- | --- | --- | --- | --- | --- | --- |
|  |  |  | Fixed 1 | Fixed 2 | Fixed 3 | R<sup>2</sup> marginal | R<sup>2</sup> conditional |
| Leaf chemistry | Anthocyanins | Climatic | PC1 | PC2 | Height 19 | 0.021 | 0.245 |
|  |  | Geographic | Site | Height 19 * |  | 0.043 | 0.258 |
|  | Chlorophyll | Climatic | PC1 | PC2 | Height 19 | 0.024 | 0.151 |
|  |  | Geographic | Site | Height 19 | - | 0.046 | 0.173 |
|  | Flavonols | Climatic | PC1 | PC2 | Height 19 *** | 0.158 | 0.277 |
|  |  | Geographic | Site * | Height 19 *** | - | 0.199 | 0.311 |
|  | Nitrogen Balanced Index | Climatic | PC1 | PC2 | Height 19 * | 0.053 | 0.185 |
|  |  | Geographic | Site | Height 19 * | - | 0.081 | 0.208 |
| Phenology | Duration bud set 2018 | Climatic | PC1 | PC2 | Cutting | 0.008 | 0.036 |
|  |  | Geographic | Site | Cutting | - | 0.022 | 0.053 |
|  | Duration bud set 2019 | Climatic | PC1 | PC2 | Cutting | 0.009 | 0.045 |
|  |  | Geographic | Site | Cutting | - | 0.020 | 0.056 |
|  | Duration bud flush 2019 | Climatic | PC1 | PC2 * | Height 18 *** | 0.058 | 0.144 |
|  |  | Geographic | Site *** | Height 18 *** |  | 0.093 | 0.171 |
|  | Duration bud flush 2020 | Climatic | PC1 | PC2 | Height 18 *** | 0.052 | 0.112 |
|  |  | Geographic | Site | Height 18 *** |  | 0.065 | 0.126 |

## 3. Results

### 3.1. Phenotypic variation between geographic regions and along climatic gradients

Phenotypic differences along climatic gradients and among ramets originating from ortets of different geographical origins were tested using linear mixed models (Table 2). Four phenotypic traits showed significant correlation with climatic gradients. The number of stems and herbivory damage (p = 0.034 and p = 0.003, respectively) were correlated with the temperature gradient (PC1), and active growth rate and duration of bud flush 2019 (p = 0.036 and p = 0.044, respectively) were correlated with the precipitation gradient (PC2). Significant differences among trees from different site of origin (Figure 1) were also found for the number of stems (p = 0.025), duration of bud flush in 2019 (p = 0.019). Autumn height in 2019, LMA, SLA, and leaf flavonol content showed a significant correlation with site of origin in the LMMs (p < 0.05 in all four traits), but no significant differences between groups after p-value correction. Supplementary figures 4 and 5 show boxplots for all traits where no significant differences were found after correcting the p-values for multiple testing.

### 3.2. Effects of common garden microenvironmental conditions on phenotypic variation

The linear mixed effects models also informed us about the respective importance of the ramets’ site of origin vs. microenvironmental conditions in the common garden (difference among blocks) on phenotypic variation(Table 2). In most traits, site of origin and climatic conditions of the ortets accounted for less than 10% of the total phenotypic variation, with the hightest fraction of the variation in tree height 2019 and 2020 (marginal R^2^ = 0.102 and 0.113, respectively) and flavonol content (R^2^ = 0.199). For all traits, conditional R^2^ values were considerably higher than marginal R^2^ values (Table 2), indicating that unknown microenvironmental variability in the common garden explained a larger fraction of the phenotypic variation among the ramets..

### 3.3. Correlation among phenotypic traits

To assess if variation in cutting diameter and developmental processes such as number of stems produced might have an effect on phenotypic traits, the relationships among phenotypic traits were assessed using Pearson’s Product-Moment Correlation (Figure S3). Unsurprisingly, most growth traits were intercorrelated. Tree height in 2018, 2019, and 2020 were positively correlated with active growth rate and negatively correlated with cutting diameter. Tree height was also correlated with ecophysiological traits (LMA and SLA), phenological traits (bud set 2018 and bud flush 2019), and traits related to leaf chemical compounds (flavonol content and NBI). Phenology and leaf chemistry traits also showed intercorrelation. The number of days that trees needed from the start of the bud flush period until the leaves were fully unfolded in 2019 (bud flush 2019 in the tables) were significantly correlated with the number of days required for a complete bud set in 2018 (bud set 2018). There was, however, no correlation between the duration of bud flush in 2020 and the duration of bud set in 2019. A few traits were found to be correlated with climatic variables from the ortet growing sites (PC1 and PC2). Cutting diameter and herbivory damage were positively and negatively correlated, respectively, with the temperature gradient (PC1). Growth rate and bud flush 2019 showed a weak negative correlation with the precipitation gradient (PC2). Bud flush 2020 and bud set 2018, on the other hand, were negatively correlated with PC2.

## 4. Discussion

In this study, we investigated the effect of previous environments on the clonal offspring of Lombardy poplar ramets growing under common garden conditions. We found that phenotypic variation of the ramets correlates with geographical and climatic gradients, but that uneven new microenvironmental conditions in the common garden might outweight parental effects.

Four of the phenotypic traits (growth rate, number of stems produced at sprouting, herbivory damage and duration of bud flush) showed a significant correlation with climatic gradients, and six traits (tree height, number of stems, LMA, SLA, flavonol content and duration of budflush) showed significant differences among ramets from different geographical regions. Both bud phenology and tree growth can directly affect tree performance and fitness (Cook et al., 2012), and have been shown to be epigenetically regulated to a certain point (Bräutigam et al., 2013, Rios et al., 2014; Lu et al., 2019). The presence of an epigenetic memory would be a major advantage for new saplings if the relevant environmental conditions remain relatively constant over time. The environmental requirements that are needed for dormancy break and bud flush are still not fully understood and are species-dependent, but it is widely accepted that temperate tree species rely on temperature and photoperiodic cues to trigger bud flush (Rohde and Bhalerao, 2007; Ibáñez et al, 2010; Malyshev et al., 2018). In poplar, after the chilling requirements and a certain day length threshold are reached, growing temperatures will trigger the processes related to bud flush (Singh et al. 2016). Consequently, under the same temperature and light conditions, no differences in the number of days that ramets from the same genotype needed for bud flush would be expected. The fact that we found significant differences in tree growth traits and bud flush, suggests that environmentally induced epigenetic memory might play a role. Although differences in tree height persisted over two growing seasons, the differences in the duration of bud flush seem to have disappeared in 2020, suggesting that the parental effects might not be very stable (Shi et al., 2019). This could potentially also be an advantage, if the survival of the clones depends on the existence of a mechanism of rapid acclimation to unpredicted conditions. For example, multiple studies have suggested that high phenotypic plasticity levels associated with epigenetic diversity might contribute to the successful establishment of clonal (and often invasive) plant species (Davidson et al., 2011; Richards et al., 2012; Mounger et al., 2020).

The phenotypic variation among ramets from different geographic origins observed in our common garden was generally low. As evidenced by the considerably low marginal R^2^ values of the linear models, in several traits this variation was not explained by the geographical origins of the ramets or by environmental gradients. Conditional R^2^ values were, in comparison, larger than marginal R^2^ values for all traits. In the models, the only variable considered as random effect was the block of the common garden where each ramet was planted. Our results indicate that microenvironmental conditions in the common garden explained a major fraction of the phenotypic variation found between ramets. The exact causes of environmental differences between blocks are unknown since we did not characterize them. However, we suspect that soil composition heterogeneity, combined with daily watering during summer droughts (Schuldt et al., 2020), could have contributed to heterogeneity in growing conditions in the common garden field. Small-scale biotic and abiotic conditions experienced by individuals have been shown to dramatically influence phenotypic plasticity, genetic variation and population persistence (Wu, 1994; Crutsinger, 2015; Denney et al., 2020). Our findings suggest that parental effects do exist and could be detected in our common garden, but they might be relatively weak compared to the micro-environmental conditions of the garden. Additionally, other traits like cutting diameter might act as confounding factors. Our results suggest that general plastic responses to extreme environmental conditions could possibly outweight parental effects, masking any potentially inherited epigenetic variation.

## Conclusions

Although intraspecific phenotypic variation in *Populus sp*. has been shown to have a large genetic component, our results indicate that the phenotypic differences found between genetically indentical ramets under common garden conditions can partially be attributed to shared environmental conditions, and might be transmitted as part of the epigenetic memory. However, we also found that uneven microenvironmental conditions in the common garden had a significant effect on the observed phenotypic variation, possibly overwriting parental effects and thus allowing for short term acclimation to new environemental conditions. In recent years, epigenetic variation has been shown to play a bigger role on phenotypic plasticity than previously thought. Further experimental research, in particular large-scale studies that combine phenotypic and epigenomic data, is necessary to understand the effects of natural epigenetic variation on phenotypic variation.

## Author contributions

Bárbara Díez Rodríguez: Methodology, formal analysis, Investigation, writing - original draft, visualization

Cristian Peña: Writing – Review & Editing

Paloma Pérez Bello: Writing – Review & Editing

Julius Bette: Investigation

Lena Lerbs: Investigation

Tabea Mackenbach: Investigation, Writing – Review & Editing

Sven Wulle: Investigation

Emanuele De Paoli: Writing – Review & Editing, Formal analysis, Supervision

Koen J.F. Verhoeven: Writing – Review & Editing, Supervision, Project administration, Funding acquisition

Katrin Heer: Conceptualization, Writing – Review & Editing, Supervision, Project administration, Funding acquisition

Lars Opgenoorth: Conceptualization, Writing – Review & Editing, Supervision, Project administration, Funding acquisition

## Acknowledgments

The authors want to thank Philipp Kurth and David Löning for their help with sampling, and Phillip Kaldeway, Leonie Braasch and Tabea Giese for their contributions to data collection. We also want to thank all the members of the EpiDiverse Consortium for their support. We are grateful to An Vanden Broeck for her advice.

## Data availability

The data that support the findings of this study are openly available in Zenodo at https://doi.org/10.5281/zenodo.5995424

