## Supplementary Information for "An uncommon garden experiment: microenvironment has stronger influence on phenotypic variation than epigenetic memory in the clonal Lombardy poplar"

### Supplementary figures and tables

### Supplementary Figure 1.

Common garden block design.

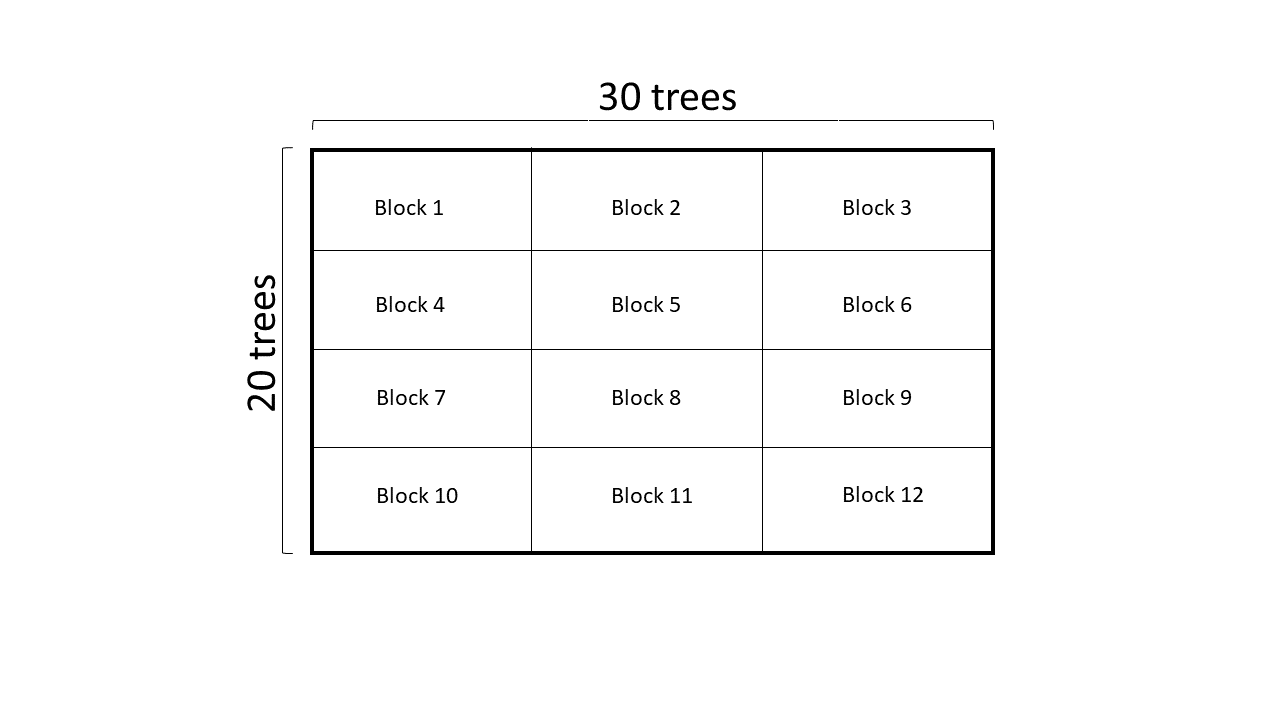

### Supplementary Figure 2.

Genotyping of Lombardy poplar ramets based on targeted genotyping-by-sequencing. Different colors indicate ramets collected in different geographic sites. Black triangles represent the three adult *P. nigra cv “Italica*” clones from the Botanical Garden in Marburg that were included in the sequencing design as controls.

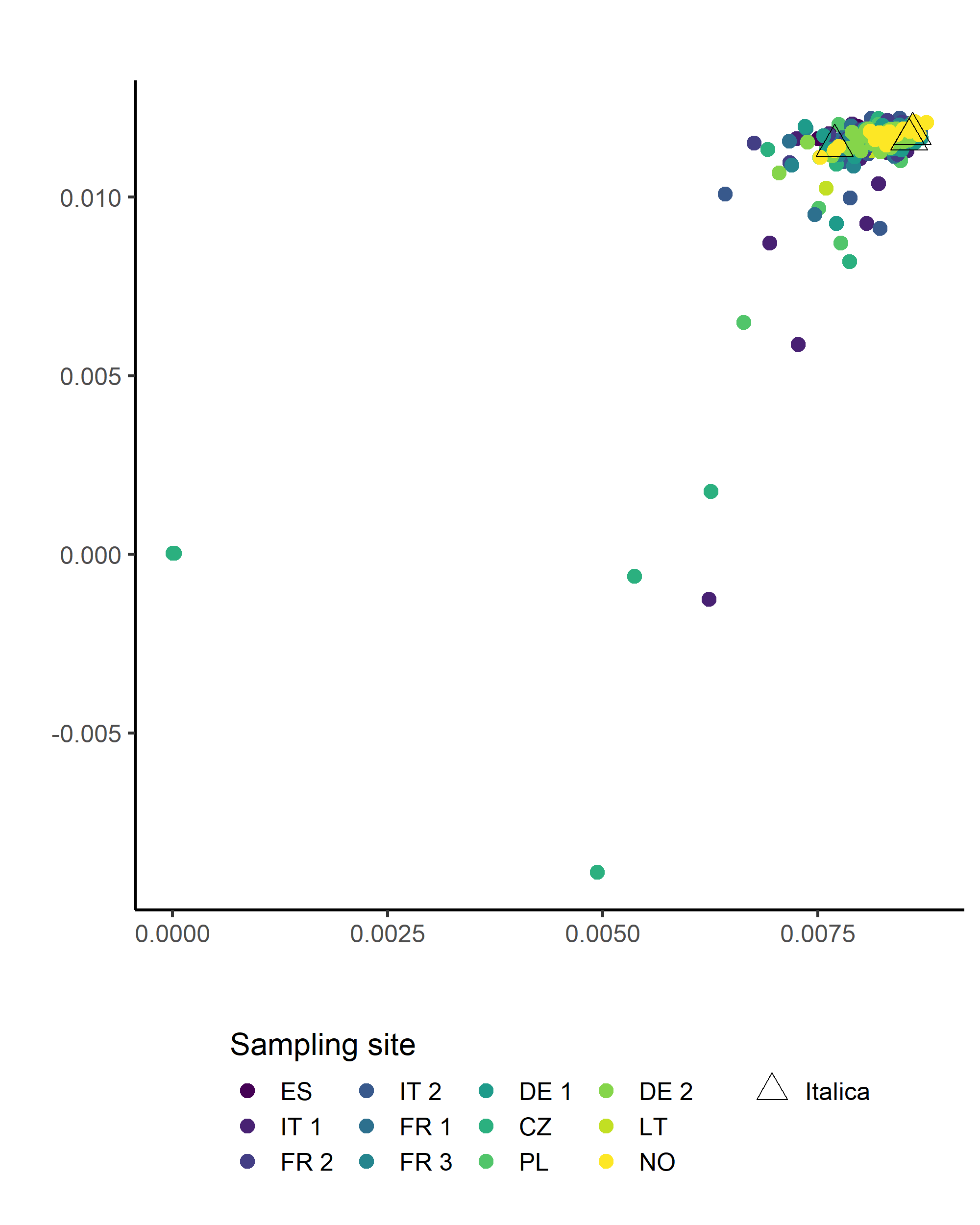

#### Supplementary Figure 3.

Phenotypic trait correlations. Pearson's Product‐Moment Correlation coefficients (*r*) indicated among all growth, ecophysiology, biotic stress, leaf chemistry and phenology traits listed in Table 1, and climatic gradients (PC1 and PC2). Colored squares correspond with the coefficients of significant correlations.

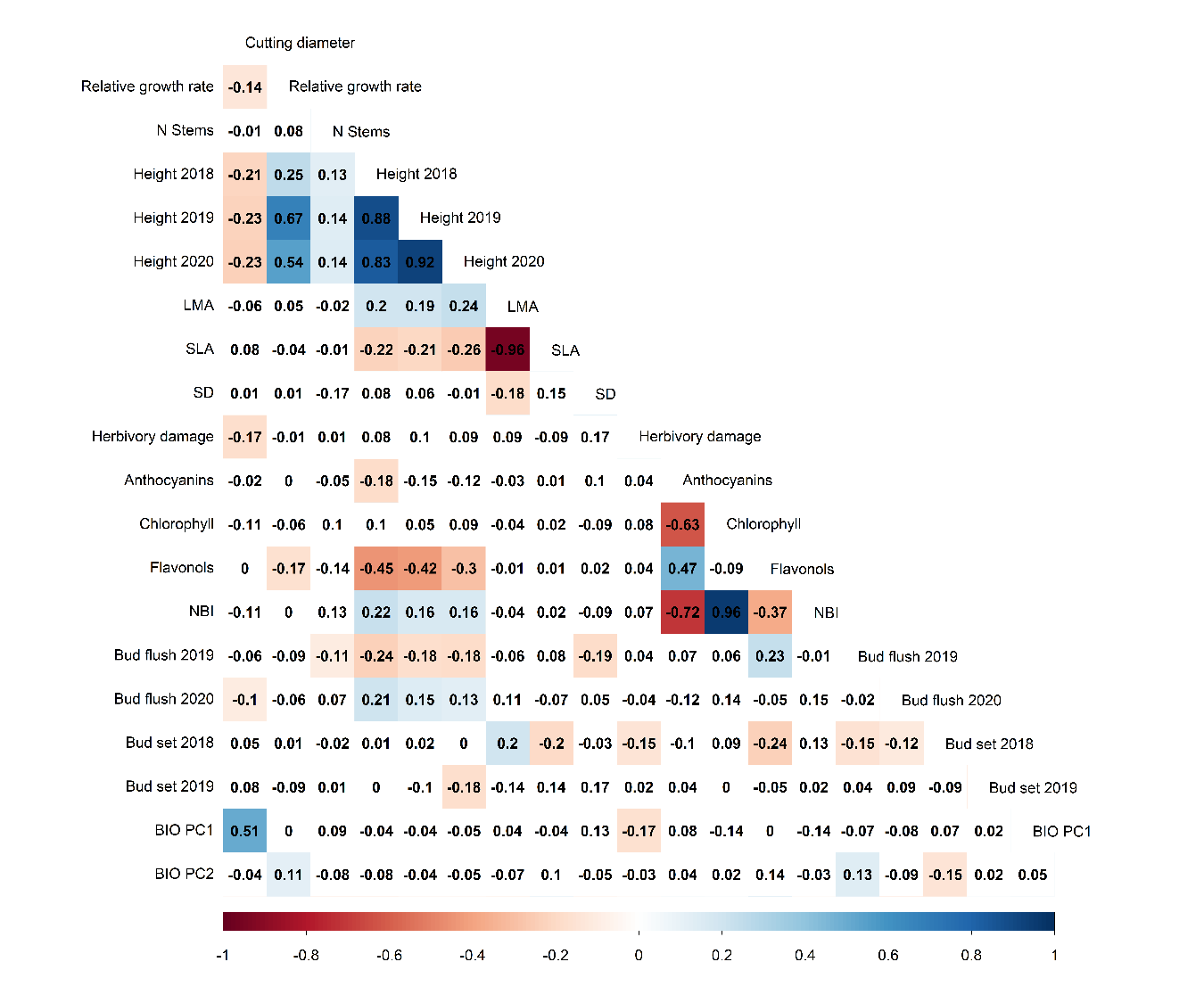

#### Supplementary Figure 4.

Phenotypic variation in traits where no significant differences between geographic origins were found. Boxplots are ordered by site, from south to north according to their geographic coordinates and labelled by the sample site code (ISO 3166 standard country code): 1. Spain; 2. Italy 1; 3. France 2; 4. Italy 2; 5. France 1; 6. France 3; 7. Germany 1; 8. Czech Republic; 9. Poland; 10. Germany 2; 11. Lithuania; 12. Norway.

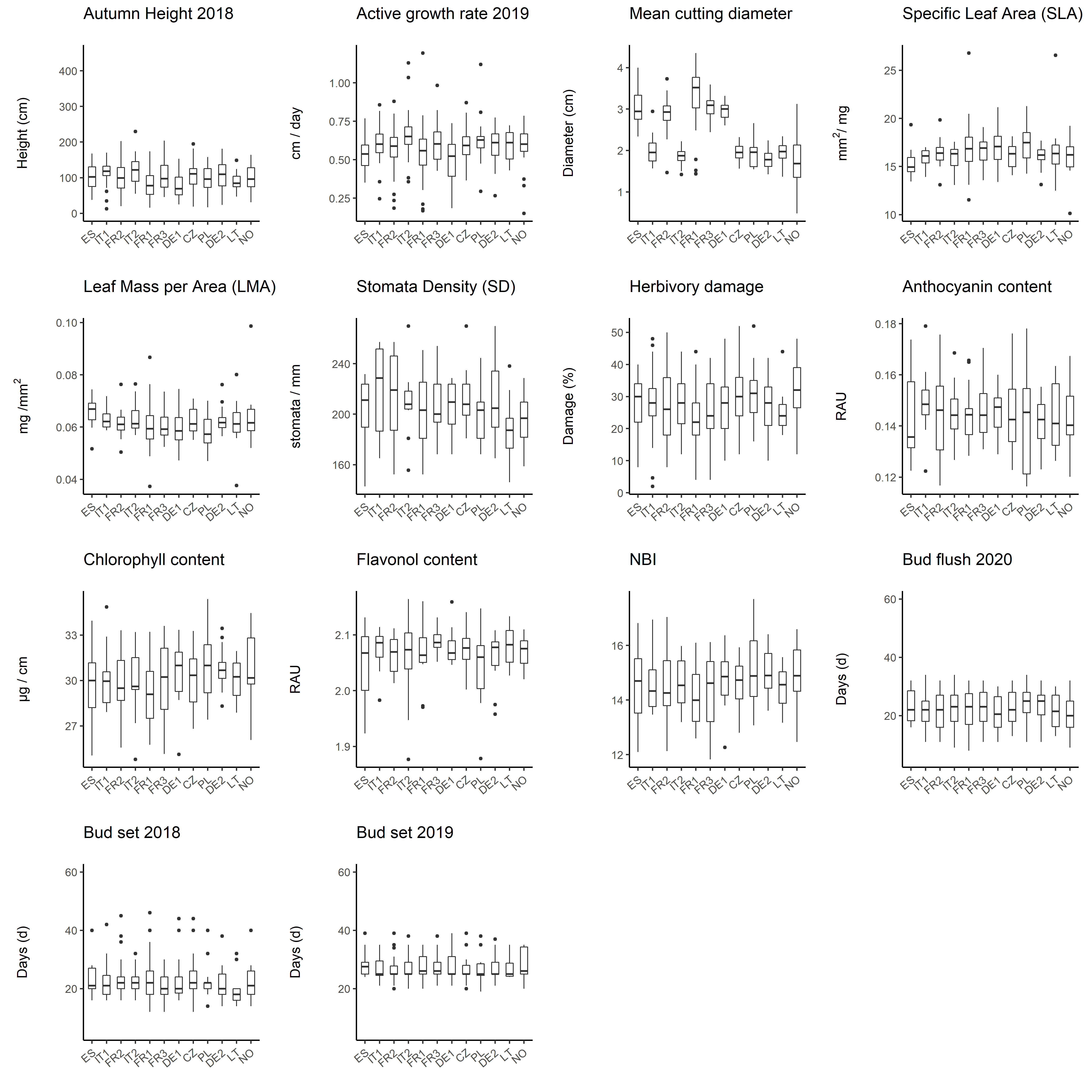

#### Supplementary Figure 5.

Phenotypic variation in traits where no significant differences between geographic origins were found. Boxplots are ordered by common garden block.

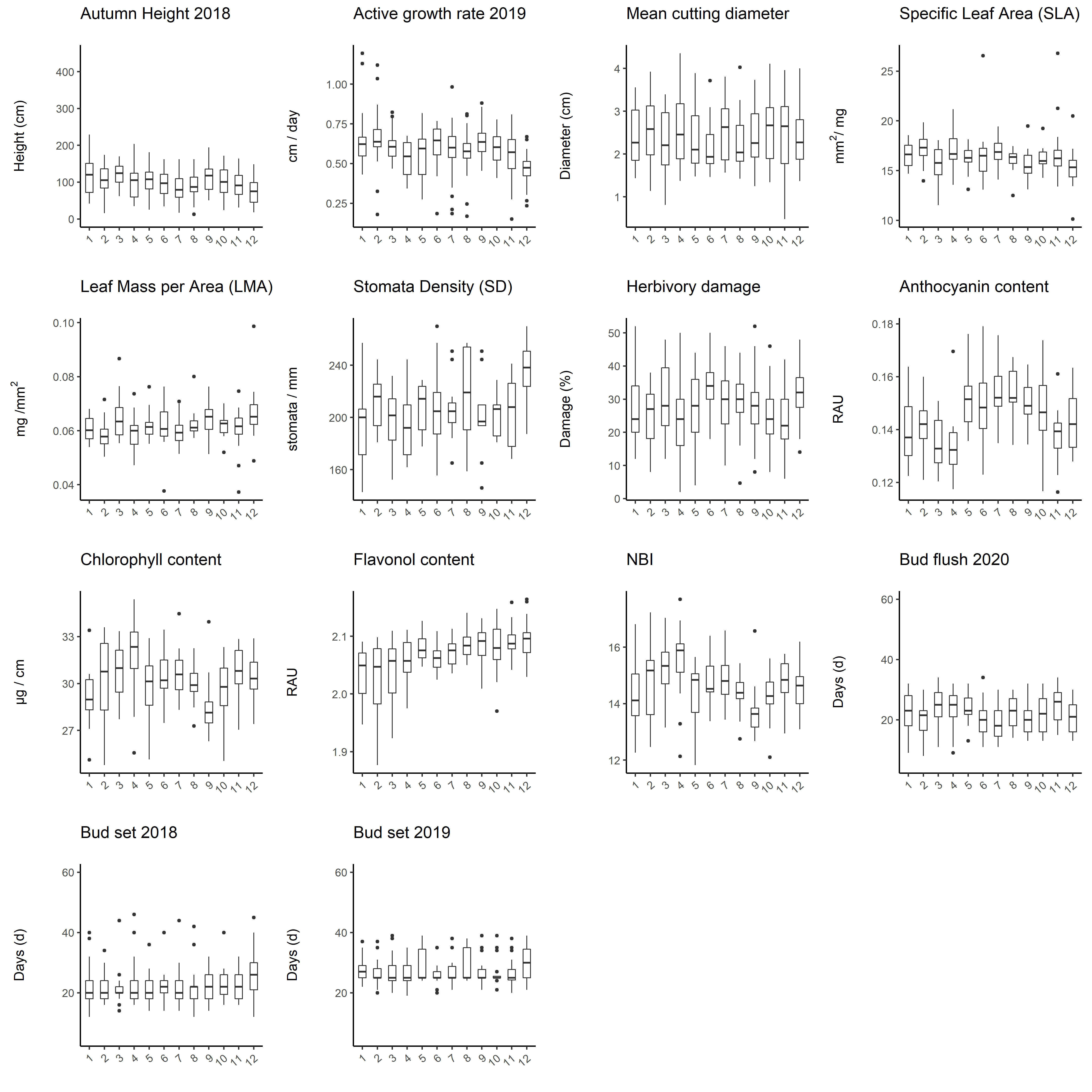

#### Supplementary Table 1.

Description and range of all bioclimatic (Bioclim) variables included in the PCA analysis, and their contributions to principal component 1 (% PC1, total variation explained = 68.1%) and Principal Component 2 (% PC2, total variation explained= 13.9%). Bioclimatic data for each of the locations of the parental clones were obtained from the CHELSA Timeseries data set.

| BIOCLIM | Description | | % PC1  (68.1 %) | % PC2  (13.9 %) | Range |
| --- | --- | --- | --- | --- | --- |
| BIO 1 | Annual Mean Temperature (°C) | | 9.31 | 2.47 | 6.42 - 15.24 |
| BIO 2 | Mean Diurnal Range (°C) | | 2.98 | 7.05 | 5.45 - 9.18 |
| BIO 3 | Isothermality (%) | | 5.48 | 2.03 | 21.32 - 33.96 |
| BIO 4 | Temperature Seasonality (°C) | | 5.38 | 0.08 | 526.51 - 790.63 |
| BIO 5 | Max Temperature of Warmest Month (°C) | | 7.36 | 4.67 | 21.02 - 30.29 |
| BIO 6 | Min Temperature of Coldest Month (°C) | | 9.50 | 1.25 | -6.64 - 4.29 |
| BIO 7 | Temperature Annual Range (°C) | | 0.99 | 2.03 | 22.82 - 29.79 |
| BIO 8 | Mean Temperature of Wettest Quarter (°C) | | 3.45 | 1.74 | 6.81 - 19.42 |
| BIO 9 | Mean Temperature of Driest Quarter (°C) | | 8.92 | 1.61 | -3.61 - 24.15 |
| BIO 10 | Mean Temperature of Warmest Quarter (°C) | | 7.55 | 3.61 | 16.67 - 24.75 |
| BIO 11 | Mean Temperature of Coldest Quarter (°C) | | 9.90 | 1.54 | -3.61 - 7.61 |
| BIO 12 | Annual Precipitation (mm) | | 4.56 | 11.82 | 391.45 - 951.67 |
| BIO 13 | Precipitation of Wettest Month (mm) | | 1.36 | 8.96 | 45.22 - 119.41 |
| BIO 14 | Precipitation of Driest Month (mm) | | 2.90 | 13.11 | 12.00 - 54.16 |
| BIO 15 | Precipitation Seasonality (%) | | 3.52 | 2.74 | 10.65 - 47.23 |
| BIO 16 | Precipitation of Wettest Quarter (mm) | | 1.49 | 7.85 | 128.46 - 332.24 |
| BIO 17 | Precipitation of Driest Quarter (mm) | | 3.76 | 12.40 | 39.00 - 172.48 |
| BIO 18 | Precipitation of Warmest Quarter (mm) | | 2.40 | 11.76 | 39.00 - 248.59 |
| BIO 19 | Precipitation of Coldest Quarter (mm) | | 9.19 | 3.11 | 63.20- 287.06 |

#### Supplementary Table 2.

List and descriptions of all traits measured under common garden conditions, and the units in which each trait was measured.

| Trait | Description |
| --- | --- |
| Tree growth |  |
| Cutting diameter (cm) | Diameter of planted cuttings |
| Height Autumn (cm) | Tree height from ground to apical shoot at the end of each growing season |
| Growth rate (cm day^-1^) | Ratio between height gain and number of days in the growth period |
| Growth period (d) | Number of days between budflush stage 5 and budset stage 2.5 |
| Stems (#) | Number of stems at the end of the growing season |
| Ecophysiology |  |
| SLA (mm² mg^-1^) | Ratio between leaf area and leaf dry weight |
| LMA (mg mm^-2^) | Inverse of SLA |
| Stomatal Density (mm^-^²) | Number of stomata per mm² |
| Biotic stress |  |
| Herbivory damage (%) | Percentage of damage caused by herbivores |
| Level of rust infection (SU) | Level of infection on a scale of 1-60, calculated from leaf-level damage and tree-level damage. |
| Leaf Chemistry |  |
| Chlorophyll (µg cm^-2^) | Chlorophyll content based on UV optical absorbance measurements |
| Flavonols (RAU) | Flavonol content based on UV optical absorbance measurements |
| Anthocyanins (RAU) | Anthocyanin content based on UV optical absorbance measurements |
| Nitrogen Balanced Index | Ratio of chlorophyll and flavonol content |
| Phenology |  |
| Budset (d) | Number of days between stage 2.5 and stage 0 of the bud set period |
| Budflush (d) | Number of days between stage 2 and stage 5 of the bud flush period |
